# Amphetamine reduces utility encoding and stabilizes neural dynamics in rat anterior cingulate cortex

**DOI:** 10.1101/2020.03.24.005256

**Authors:** Saeedeh Hashemnia, David R. Euston, Aaron J. Gruber

**Affiliations:** Canadian Center for Behavioral Neuroscience, Dept. of Neuroscience, University of Lethbridge, Lethbridge, AB, Canada

**Keywords:** Ensemble recording, choice, effort, state space, neuromodulation

## Abstract

The anterior cingulate cortex (ACC) appears to support decisions by encoding the effort-reward utility of choice options. We show here that d-amphetamine (AMPH) has dose-dependent effects on this encoding and on neural dynamics in rat ACC that are concordant with its behavioral effects. Low-dose AMPH increased task engagement and had mild effects on neural encoding, whereas high doses disrupted utility signaling and decreased task engagement. The disruption involved reduced reward signaling and compressed effort-reward encoding of utility cells, which corresponded with reduced reward consumption behaviors. Furthermore, low-dose AMPH stabilized and accelerated trajectories of neural activity in state-space, whereas high-dose AMPH destabilized trajectories. We propose that low-dose AMPH increases both excitability and stability, which preserves information and accelerates evolution of a neural ‘script’ for task execution. Excessive excitability at high doses overcomes stability enhancement to suppress weakly encoded features (e.g. reward) and cause deviation from the script, which interrupts task performance.

**Significance Statement:** Amphetamine reduced reward signaling by individual neurons in rat prefrontal cortex, but increased the stability of ensemble dynamics. These effects account for animals’ increased task engagement, despite reduced reward intake.

## Introduction

Rats typically choose to exert increased effort, such as barrier climbing or lever pressing, if the associated reward is of considerably higher value than that of lower-effort choice options. This preference is reduced or eliminated by lesions of the Anterior Cingulate Cortex (ACC) (Walton *et al*., 2002; Walton *et al*., 2003; Schweimer & Hauber, 2005; Holec *et al*., 2014), lesions of ventral striatum (VS) (Hauber & Sommer, 2009), or by disruption of dopamine transmission in these areas (Cousins *et al*., 1996; Schweimer & Hauber, 2006; Mai *et al*., 2012). Therefore, drugs such as d-amphetamine (AMPH) that increase extracellular dopamine levels in these structures (Chiueh & Moore, 1973; Pum *et al*., 2007), will likely influence effort-reward choices. Indeed, choice preference is biased toward high-effort, high-reward options by systemic AMPH administration (Floresco *et al*., 2008; Bardgett *et al*., 2009). It remains unclear what aspects of neural information processing are affected to produce this effect.

Effort, reward, and other features pertinent for economic decisions are often formalized within the concept of utility (Phillips *et al*., 2007). Options requiring low effort but yielding a large food reward have high utility to hungry animals, whereas options requiring high effort or yielding little/unwanted food have low utility. Because utility can be expressed as the expected value discounted by its associated costs, the bias toward high-effort high-reward options under AMPH could reflect a reduction (or ‘discounting’) of effort, or an increase in expected reward value. Behavioral data, however, suggest that reward value is *decreased*. AMPH-treated rats appear less motivated to eat; their latency to first consumption is longer, they consume less than usual, and they spend less time eating (Blundell *et al*., 1976; Blundell *et al*., 1979; Leibowitz *et al*., 1986) . Suppression of food-intake by AMPH is also evident in primates (Foltin, 2001). Even though AMPH appears to decrease reward value, it induces rodents and primates to become more engaged in learned food-seeking behaviors (Foltin, 2001; Odum & Shahan, 2004). This suggests that they are more willing to work for the reward, as if the food had higher utility, but then show less interest in consuming the reward, as if the food had lower utility. Alternately, the apparent shift in utility could be a by-product of increased motoric output {Shin, 2010 #71}, but this does not explain why animals engage in a task for an unwanted reward, rather than some other behavior. The behavioural data, therefore, do not support a clear prediction about how AMPH may influence the neural encoding of utility.

Electrophysiological recordings indicate that the ACC and nearby regions in the medial prefrontal cortex (mPFC) encode a variety of signals related to choice. These include the position of the animal (Euston & McNaughton, 2006; Fujisawa *et al*., 2008; Mashhoori *et al*., 2018), task phase (Lapish *et al*., 2008; Balaguer-Ballester *et al*., 2011), reward (Gruber *et al*., 2010; Cowen *et al*., 2012), choice (Cowen *et al*., 2012), effort (Cowen *et al*., 2012; Hashemniayetorshizi *et al*., 2015), and other task-related features (Cowen & McNaughton, 2007; Gruber *et al*., 2009; Durstewitz *et al*., 2010; Sul *et al*., 2010). These are consistent with findings in monkeys and humans (Isomura *et al*., 2003; Kennerley *et al*., 2006; Croxson *et al*., 2009; Skvortsova *et al*., 2014; Blanchard *et al*., 2015; Klein-Flugge *et al*., 2016). Neural recordings have shown encoding of costs-benefit computations in rat ACC (Hillman & Bilkey, 2010; Cowen *et al*., 2012). Moreover, AMPH modulates neural dynamics of ACC activity (Lapish *et al*., 2015), but this has not been explicitly shown in the effort-reward encoding of neurons. Here, we attempt to link these independent observations to better understand how systemic AMPH affects the encoding of utility by single units and ensembles in the ACC. We propose that reduced reward interest is linked to decreased reward signaling by AMPH, but that animals continue to engage in the task because ensemble neural dynamics become ‘trapped’ in task-related trajectories.

## Material and Methods

### Subjects and surgical procedure

In this study, four adult male Fischer Brown Norway (FBN) hybrid aged 6 to 10 months were used. Rats were born and raised on-site, housed individually in a 12h-12h reverse light cycle, and habituated to handling for two weeks prior to surgery. The fabrication and surgical implantation of head-mounted drives was completed as previously described (Euston & McNaughton, 2006), but is explained briefly here. Surgeries were carried out prior to any training. Animals were deeply anesthetized with isoflurane throughout the procedure (1-1.5 % by volume in oxygen at a flow rate of 1.5 L/min). Each animal was implanted with a “hyperdrive” consisting of 12 independently-movable tetrodes (McNaughton *et al*., 1983; Wilson & McNaughton, 1993) and 2 reference electrodes. The hyperdrive bundle was centered at 3.00 mm AP, and 1.3 mm ML of left mPFC and angled 9.5 degrees toward the midline. A craniotomy was made around the electrode exit site of the drive, and the dive bundle was lowered to the brain surface. The dura was retracted, and the hyperdrive body was secured to the skull with anchor screws embedded in dental acrylic (Lang Dental, Wheeling, US). Following surgery, rats were administered daily injections of 1mg/kg Metacam (analgesic) for 3 days and 10 mg/kg Baytril (antibiotic) for 5 days. Tetrodes were lowered 950 μm from the skull surface after the surgery and then gradually lowered over the next 2-3 weeks to reach the target depth. Food restriction started after the animal recovered from the surgery (7 days) and was monitored daily to ensure the weight was at least 85% of the free-feeding weight for the duration of the experiment. The experiments were performed in dim light during animal’s waking phase. All procedures were performed in accordance with the Canadian Council of Animal Care and the Animal Welfare Committee at the University of Lethbridge.

### Experiment

The behavioural apparatus and the method of data collection used in this study were described previously (Mashhoori *et al*., 2018). Briefly, we used an automated figure-8 maze (Fig. 1A), which is a modified version of the classic T-maze frequently used in studies of effort-reward decision-making (Salamone *et al*., 1994; Walton *et al*., 2002). The track of the maze was 15 cm wide, and configured into a rectangular pathway measuring 102 x 114cm. The maze contains a central feeder on the T-stem from which a trial was initiated. Two side feeder wells were located on two platforms located in the upper corners of the maze. The platforms could move vertically to require a variable height climb to reach the feeder. Animals descend from the platform by a ramp to return to the starting feeder. The elevation of the platform was 0 cm for the low-effort condition (level with the track) and was 23 cm for the high-effort condition. The reward was Ensure beverage (chocolate flavoured), and the volume was 0.03ml for low-reward, and 0.12 ml for high-reward. The same small volume of Ensure was delivered at the center feeder in all trials so as to motivate the rat to return to the start position, and to serve as a control. Four gates were located on the entry points of the T-stem and T-arms to prevent animals going backward. These were also used on some trials to force animals to select one target feeder. On other trials, animals were free to choose either target feeder. Over a course of roughly three weeks, animals were trained on the maze with a mixture of force and free trials. Animals were identified as well-trained when they choose high-reward options over the low-reward ones more than 80% of the trials in which the barrier height was the same on both sides. In the following sessions, the animals were required to perform two full blocks of forced trials as shown in Figure 1B. Each block of trials consisted of five groups of 20 forced trials where the effort-reward conditions were constant. Each of these trial groups was designed to investigate one of the three decision features (effort, reward, path), which was conducted by manipulating only one of these and keeping the other two constant. However, only effort and reward groups were used in this study. Trials were arranged alternatively left and right throughout the task. One of the recorded rats used a different task design, consisting of either four or six groups of 16 trials (10 alternate-side force followed by 6 free trials) in which another level of effort was also added (23 cm for medium-effort, and 46 cm for high-effort).

**Figure 1.**
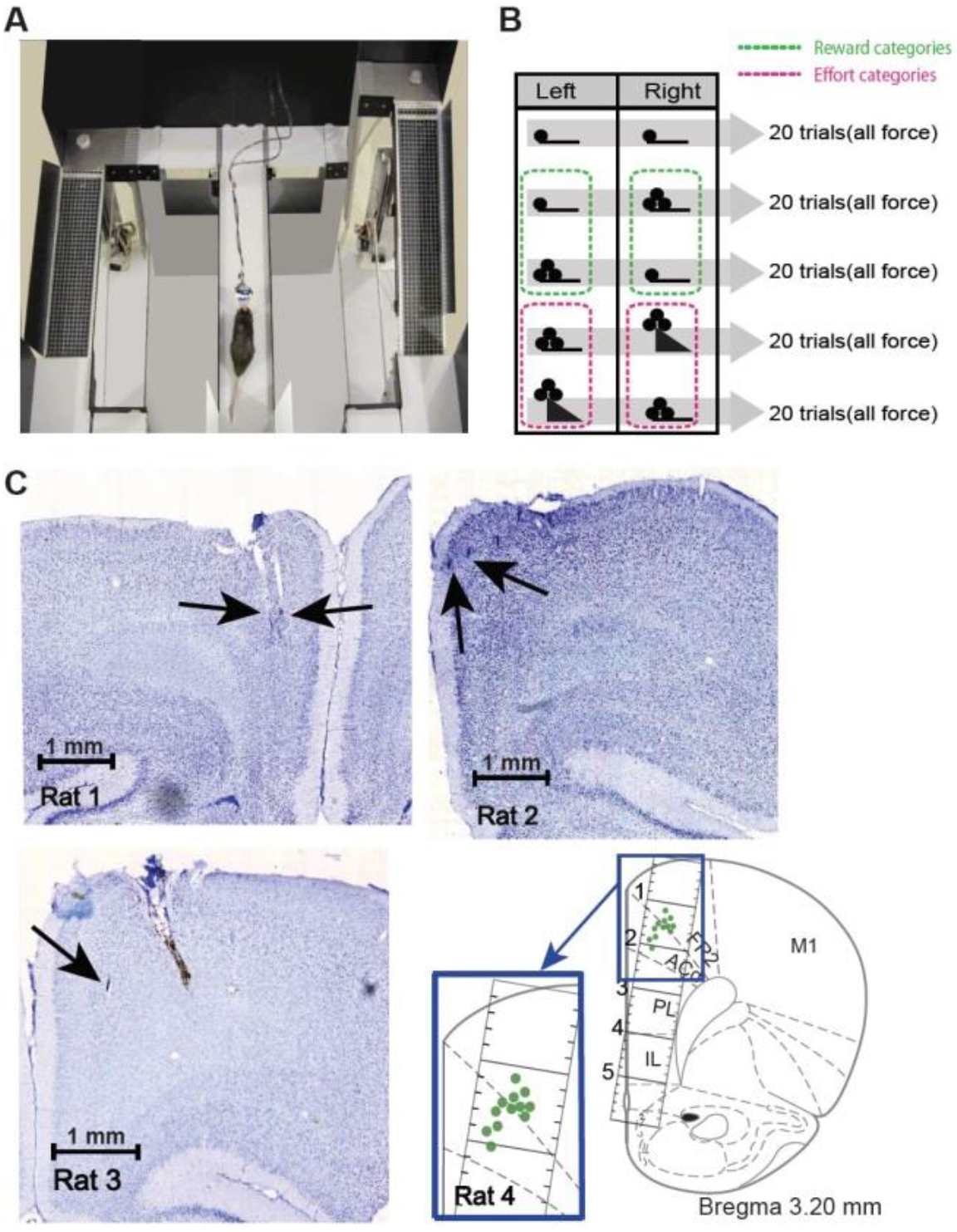
Experimental apparatus and task. A, Photograph of the figure-8 maze. The rat in the picture is consuming reward at the central feeder and the right-side platform is elevated. Side feeder wells (white circles) are located on top of the platforms, and wire-mesh ramps lead back to the starting feeder. B, Task schematic illustrating one full block of the task. The table shows the order of grouped trials in each effort-reward condition. The height of triangles indicates the barrier height and the number of filled circles represents the amount of food reward at the side feeder. The same trial sequence is repeated two times per session. Dashed lines show the subset of trials used for testing the effort (red) and reward (green) encoding. C, Histological brain sections showing endpoints locations of electrodes. The electrode marks pointed by arrows for three rats are shown on stained coronal brain slices. The estimated electrode location for another rat is also shown superimposed on a figure adapted from a stereotaxic atlas (Paxinos & Watson, 2014).

All rats were administered with saline or AMPH during a 5-10 minutes interval after finishing the first block of the experiment, before starting the second block. Three doses of AMPH (0.5, 1, 1.5 mg/kg) were used in this study. Only one dose (or saline) was given per daily session. The order of drug/saline injection across sessions was: saline, AMPH 0.5 mg/kg, AMPH 1 mg/kg, saline, AMPH 1.5 mg/kg. The rat performing the alternate task design received the saline and drugs in different orders: saline, AMPH 0.5 mg/kg, AMPH 1.5 mg/kg, saline, AMPH 1 mg/kg, saline, AMPH 0.5 mg/kg.

### Histology

Following completion of the study and 2-5 days before transcardial perfusion, recoding sites were marked by passing 10 uA direct current for 10 second through one electrode of each tetrode. Then, rats received lethal injections of sodium pentobarbital (100 mg/kg i.p.) and were perfused with PBS and 4% paraformaldehyde (PFA). The brains were post-fixed for 24 hours in 4% PFA and then transferred and stored in 30% sucrose and PBS solution with sodium azide (0.02%). After at least 24 hours, the brains were coronally sectioned at 40 μm thickness using a CM3050 S freezing Cryostat (Leica, Germany) and mounted on glass microscope slides, then stained by cresyl violet. Digital images of the prepared brain sections were produced with a Nano-Zoomer slide scanner (Hamamatsu, Japan) and visually inspected to determine the location of marking lesions. As shown in Figure 1C, electrode marks were identified for three brains. The 4^th^ brain was destroyed during histology; the estimated locations of the electrodes were determined by the daily logs of electrode depth, and are indicated in the Figure.

### Analysis

A total of 22 sessions were included in the analysis. The data include at least two sessions with saline treatment and one session of each drug dosage for each animal. In total, 1,266 cells were recorded, out of which 1,209 were assumed as pyramidal neurons and included in the analysis. Each session is partitioned into two phases, pre- and post-injection, and the analysis contrasts the relative change in the post-injection phase with respect to the pre-injection phase.

#### Preprocessing

Camera-based tracking of LEDs located on preamplifiers attached to the recording drives were used to estimate the position of animals on the maze. We used image registration methods in MATLAB to correct any maze or camera shifts or rotations between the recorded sessions in order to normalize maze position across sessions.

Recorded spikes waveforms were first automatically clustered using KlustaKwik (author: K. D. Harris, Rutgers-Newark) and then manually sorted using MClust (David Redish, University of Minnesota, Minneapolis). Putative individual neurons were then manually classified to pyramidal and interneurons based on half-amplitude and trough-to-peak durations (Bartho *et al*., 2004). Neural analysis was restricted to only putative pyramidal neurons unless otherwise stated.

#### Behavioural analysis

The effect of AMPH on task performance was investigated using four behavioural measures: the smoothness of rat locomotion trajectories; the median velocity of each trial; the number of off-task distractions; and time spent on reward consumption. The smoothness was measured by the Hausdorff (fractal) dimension (Hausdorff, 1919; Gneiting *et al*., 2012), which quantifies the roughness of an object. The object in our case is the rat’s trajectory of movement. Hausdorff dimension increases for the trajectories with movement deviating from the linear axis of the track. We first made binarized images of the running trajectory for each trial of the task. The images were 1 for any pixel the rat traversed, and 0 otherwise. These images were then superimposed (logical OR operation for each pixel over the image stack) to create a grand image of all pre-injection trials and a grand image of all post-injection trials. The dimension was next obtained for each grand image separately. In case the number of trials was not the same for the pre- and post-injection phases, we randomly sub-sampled the trials (100 times), computed Hausdorff dimension for each, and then took an average so as to ensure that this is not an effect of heterogeneous sample sizes among conditions. Secondly, running velocity was calculated by the change in rat’s position (in pixels) between every 111 msec time bin (1 pixel/sec is equivalent to 2.36 mm/sec). Thirdly, trials with off-task behaviours were found by manually scoring the video. Off-task behaviours included long pauses on the track, continuous repetition of circling, backtracking, or grooming. Since these stereotyped behaviours are known to be induced by AMPH (Randrup *et al*., 1963; Randrup & Munkvad, 1967), we expected to observe them more often in the post-injection phase. Finally, the reward consumption time was measured by the duration that the animal’s velocity dropped below a certain threshold (59 and 118 mm/sec on central and side feeders, respectively) at each of the three reward zones. The relative change in all these four measures were found with respect to the pre-injection phase, and the mean were compared statistically between saline and drug sessions (ANOVA for mean and Kruskal-Wallis for median tests)

#### Modulation of ACC single neuron activity by effort or reward

We first linearized the track by dividing each loop of the figure-8 maze into 36 spatial 2D bins, starting at the central feeder (Fig.3-inset). The firing rate of individual neurons was computed as the number of spikes in a 0.3 sec time bin centered on each of these spatial bins. In each bin, the change in firing rate was computed among trials in which only one parameter was different (trial organization shown in Fig.1B). For instance, to compute the effect of effort on neural signaling, we made conditional means from trials in which reward size and reward location were the same. A neuron was considered to be responsive to effort if it significantly discriminated low- and high-conditions in at least one of the two effort groups shown in Figure 1B (t-test or ANOVA, significant at p<0.05). The percentage of effort-responsive neurons was found in each of the 36 bins after removing the cells with no discrimination of effort, reward, or feeder location from the population. Grouping the sessions with similar treatments, the average portion of responsive neurons were obtained for pre- and post-injection phases separately. The same procedure was followed to compute sensitivity to reward.

The main aim of this study was to investigate the effect of AMPH on the encoding of cost and benefit, including the joint encoding in cells signaling ‘utility’. To address this, we first measured the bias strength of individual neurons toward effort and reward, and then compared population variability in pre- and post-injection phases. Bias was computed by the Pearson correlation coefficient between the spiking rate of each neuron and the two levels of effort or reward in each of the first 16 spatiotemporal bins (from the central feeder to the bin after the side feeder). These coefficients were then collapsed across the sessions with similar treatment. They were then plotted in a symmetric plane with axes of correlation for effort and for reward. In order to identify where the majority of neurons lie in the joint-encoding plane of effort (E) and reward (R), a polar histogram with three equal bins in each quadrant was used. Note that the plane makes four quadrants (Fig.4B-inset), which we numbered as in plane geometry, Q_I_ (0-90°): E+/R+, Q_II_ (90-180°): E+/R-, Q_III_ (180-270°): E-/R-, Q_IV_ (270-360°): E-/R+. The ‘+’ indicates positive correlation with either reward (R) or effort (E), while the ‘-’ indicates negative correlation. Next, we quantified the direction in which neurons are tuned in E-R space. We used principal component analysis (PCA), to determine the axis of maximum variability of all cells in the E-R space. We bootstrapped data by randomly selecting the same number of neurons for pre and post-injection phases for 100 times per spatial bin and found principal components (PCs) to be stable, indicating that this is a reliable method. Next, we statistically tested the difference of principal component coefficients of the two phases for each treatment (Kuiper two-sample test which is the circular analogue of the Kolmogrove-Smirnov test). Also, we tested the maximum variance explained by the first principal component and plotted the relative difference between pre- and post-injection of this parameter in each treatment condition. This plot is again based on 100-times bootstrapped data points.

#### Analysis of ACC population activity

Spikes from each trial were time-warped in each of the four important task epochs: reward-consumption at central feeder, interval from the central feeder to the barrier, climbing the barrier, and reward-consumption at target feeder. In other words, spikes were linearly expanded or contracted to have the same time-length epochs as the reference. The reference was obtained by making an average over the time spent in each of these epochs by the animals before administration of drug or saline. This reference is then used for all animals. Having the same time-length enables us to align the epochs and compare them either trial by trial or on average between groups of trials. Moreover, it controls for variability that may occur due to altered movement speed or pauses in some trials. The aforementioned task epochs were found using animal position as well as velocity with certain thresholds as explained in behavioural section. These epochs were color coded as in the insets of Figures 6 to 9.

In order to investigate the modulation of ACC population activity by effort and reward, we first used Gaussian Process Factor Analysis (GPFA) to reduce the dimensionality (Yu *et al*., 2009). GPFA extracts the latent structure embedded in the recorded population. In each experimental session, first 8 most important latent factors were extracted from the pre-injection data. These factors represent the directions in the high-dimensional space along which projections show largest variance. We removed trials in which it took the animal more than 20 sec to return back to the central from the side feeder. The post-injection trajectories were projected on the same low-dimension space in order to compare them with the pre-injection phase. That is, all trajectories from the same session are mapped into the same latent space. Finally, we represented the neural trajectories in two different ways for a visual comparison between conditions: 1) in the 3D space made by the first three GPFA factors and 2) in the 2D space of each factor versus time. Note that trials with off-task behaviours are removed from the illustrations; a separate illustration (Figure 7) shows typical examples of neural trajectories during off-task behavior. In order to decode information from the ensemble, the trajectories of the first and second latent factors were averaged over each task epoch and statistically compared by ANOVA between low- and high-effort or reward trials. Following this test, the fractions of sessions with similar treatments that significantly discriminate these features in each epoch were found for the pre- and post-injection phases.

In the next step, we computed the state-space occupancy of neural trajectories in 3D space made by the first three GPFA factors. The boundary of the 3D space was obtained by the convex hull of the trajectory from the central to target feeders; then, the enclosed volume was calculated by subdividing the object into smaller pieces using triangulation and adding them together. We computed mean volumes over each condition of interest (inject, reward volume, effort), and tested for differences in group means by RM-ANOVA. Next, the velocity of neural trajectories (in the original neural space rather than latent space) was assessed by computing the ‘ decorrelation function’, which quantifies the similarity of population activity as a function of time or position (McNaughton, 1998; Battaglia *et al*., 2004). We aimed to test if AMPH affected the speed by which population correlation of spatial firing rate decays with distance. Smaller spatial bins (200 bins total) were used to improve resolution. The place field profile of each cell was obtained by averaging the firing rate of each cell in each spatial bin. Next, the population vector cross-correlation matrix was built by calculating the matrix of correlation coefficients of all neurons as a function of spatial lag. The decorrelation function was calculated by averaging over the parallel (diagonal) elements. To quantify the decay of correlation over the distance, we found the spatial distance where this function first drops below the half-amplitude (i.e. threshold = 0.5). The relative change in the half-amplitude crossing distance of the post-injection phase with respect to the first phase was tested between saline and AMPH doses (ANOVA).

Data and analysis code are available online (https://github.com/SaeedehUleth/AMPH-and-utility-encoding)

## Results

We recorded ensembles of single neuron activity from the ACC of well-trained rats (n=4) performing a forced-alternation task in which the reward volume and the effort (barrier climbing) to enter either of two reward zones were manipulated independently. The yield was 55±6.7 (mean ± SD) simultaneously recorded cells per session, for a total 1,209 putative pyramidal cells in the analyzed dataset. We examined the effect of d-amphetamine (AMPH) on behaviour and its neural correlates by means of a pre/post design in which the drug or vehicle was administered (i.p.) halfway through the session.

### AMPH increases running speed, but decreases reward consumption and running trajectory smoothness

The running trajectory of the rats, estimated by video tracking of head-mounted LEDs, showed decreasing path smoothness as the dose of AMPH increased (Fig.2A). We quantified this by the Hausdorff fractal dimension, which assesses the distance between spatial data points in order to provide a measure of path roughness (Hausdorff, 1919; Gneiting *et al*., 2012) (Fig.2B; ANOVA, Main effect of dose on fractal dimension F_0.014,0.024_ = 3.47; p = 0.038). The median running velocity increased following administration of higher doses (1.0 & 1.5 mg/kg) as compared to saline (Fig.2C; Kruskal-Wallis, all pairwise comparisons except for saline and dose 0.5 mg/kg are significant, Chi-sq = 109.15; p = 2× 10^−23^). The effect appeared to peak at the 1.0mg/kg dose. The amount of off-task behaviour (circling, pausing, backtracking) increased with AMPH dose (Fig.2D; ANOVA, F_262.9,469.8_ = 3.36; p = 0.042), whereas the amount of time at the three reward feeders strongly decreased with increasing dose (Fig.2E; Kruskal-Wallis median test, Chi-sq = 208.9; p = 5×10^−45^; df = 3 for central feeder, Chi-sq = 148.76, p = 4×10^−32^; df = 3 for low-reward feeder, Chi-sq = 136.24; p = 2 × 10^−29^; df = 3 for high-reward feeder). Post-hoc tests of means revealed that all pairwise comparisons were significantly different (p<0.00001), except between 1 and 1.5 mg/kg, and between saline and 0.5 mg/kg for the side feeders.

**Figure 2.**
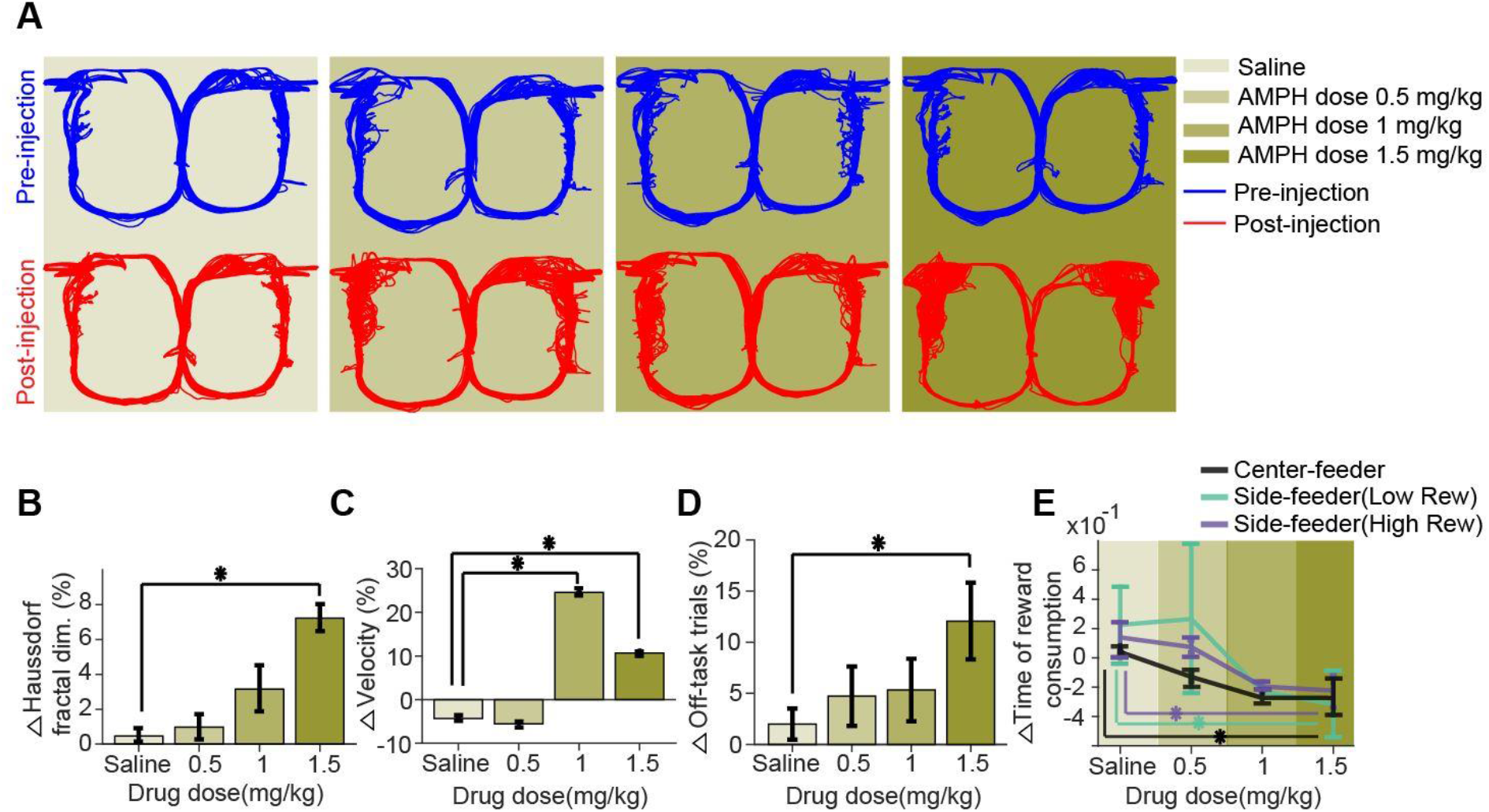
Effects of AMPH on task performance. A, Representative examples of locomotion trajectories superimposed for all trials of each session for one rat before or after injection of saline or AMPH. B, Mean change of Hausdorff fractal dimension after injection of saline/AMPH for all rats, showing an increase in path variance with increasing AMPH. C, Median change of velocity. D, Mean change in the relative proportion of trials with off-task behaviour. E, Median change in the time of reward consumption. Error bars show the Standard Error of the Mean (SEM) or Median (SEMd) (*: statistically significant at p<0.05 - but only comparisons with saline are illustrated).

These data indicate that engagement in the task decreases with higher doses of AMPH, while lower concentrations do not appear to affect it significantly. Moreover, rats did not complete a sufficient number of trials for analysis following administration of 2.0mg/kg (not shown), further supporting this observation. These data indicate that AMPH increases motoric output, but decreases feeder engagement, which is consistent with previous reports (Randrup *et al*., 1963; Randrup & Munkvad, 1967; Blundell *et al*., 1976; Blundell *et al*., 1979; Leibowitz *et al*., 1986; Floresco *et al*., 2008).

### AMPH does not affect the proportion of effort- or reward-responsive ACC neurons

We next examined if AMPH administration affected single unit encoding of effort or reward in a manner that may explain its behavioural effects. We first linearized the maze by partitioning the track into 36 two-dimensional bins in order to facilitate analysis of neural signaling during specific epochs of the task (Fig.3A inset). The percentage of recorded neurons discriminating the ramp height (i.e. encoding effort) gradually elevates from about 10% at the central feeder to a peak near 35% at the barrier, then sharply drops after climbing (Fig.3A). The percentage of cells discriminating effort again increases to about 25% while the animal is descending the ramp back to the central feeder. A smaller proportion of ACC neurons (10-20%) encode reward volume (Fig.3B), consistent with previous reports (Cowen *et al*., 2012). The apparent change in the smoothness of these plots at high doses is most likely an artifact of sampling due to fewer neurons recorded in these sessions. The relative difference between the proportion of effort (and reward) selective neurons before and after the injection is not significantly different across drug doses and saline (Fig.3; ANOVA p>0.05 in all spatial bins). Thus, AMPH treatment does not appear to affect the proportion of cells encoding effort or reward. It is possible, however, that AMPH affects the firing rate of these cells. Therefore, we next investigated how the encoding of these variables by single units is affected by AMPH.

**Figure 3.**
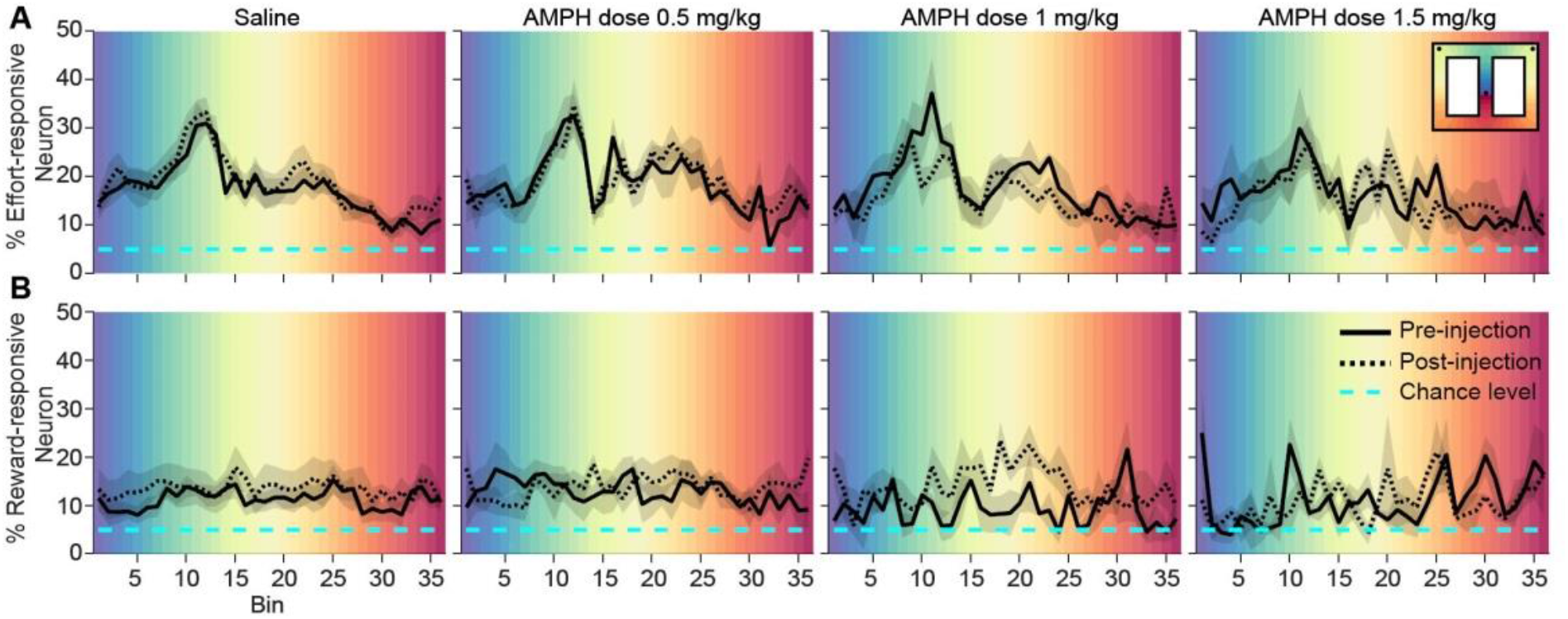
The proportion of ACC neurons encoding effort or reward. A, Mean proportion of recorded neurons that discriminate barrier height in pre- or post-injection conditions. B, Mean proportion of recorded neurons responding to the reward in pre- or post-injection conditions. The shaded region surrounding the curves indicates SEM. Background colors correspond to each of the 36 spatial bins of the maze shown in the inset.

### AMPH compresses the encoding of utility by single-units

The analysis of costs and benefits is typically formalized through the concept of utility, which can include many features (Kennerley *et al*., 2006; Hayden *et al*., 2009; Kennerley *et al*., 2009; Kennerley & Wallis, 2009; Hillman & Bilkey, 2010; Amemori & Graybiel, 2012; Porter *et al*., 2019). Here, we focus on the joint encoding of effort and reward by ACC neurons. The analysis described above revealed that the largest proportion of effort-encoding neurons modulate their activity in the several bins prior to the barrier climb. We, therefore, analyzed neural activity during this task phase. We computed the correlation of each neuron’s firing rate with effort, and independently computed its correlation with reward, in each spatial bin from the central feeder to the side feeders. A scatter plot of these correlation coefficients before drug administration suggests that the encoding of reward and effort among individual neurons are anti-correlated (Fig.4A). We quantified the relationship by the slope of the first principal component (PC) of the effort-reward correlation data, which is negative, indicating that the maximum variance of data is explained by neurons located in the second (Q_II_) and fourth (Q_IV_) quadrants. Figure 4B shows the distribution of neurons in the effort-reward coding plane at four locations of the maze. Neurons of quadrants Q_II_ and Q_IV_ have opposing signaling of utility. Neurons of Q_IV_ tend to generate more action potentials for high-utility conditions (i.e. when the effort is low or the reward volume is high), and fire less in low-utility conditions (i.e. high-effort or low-reward). Neurons in Q_II_ exhibit the inverse relationship among firing and value. The slope of the first PC becomes steeper with increasing AMPH (Fig.4C), indicating a loss of neural firing correlation with reward. Circular statistical analysis reveals that the post-injection PCs are significantly different than the pre-drug condition for AMPH, whereas saline injection has no effect (Kuiper two-sample test, test statistic k = 160; p = 0.028 for 0.5mg/kg, k = 176; p = 0.007 for 1mg/kg, k = 240; p < 10^−3^ for 1.5mg/kg AMPH). To ensure that this is not an effect of heterogeneous sample sizes among conditions, we ran a bootstrap analysis (100 repetitions) in which we randomly sub-sampled the data to obtain the same number of neurons for each condition. The PCs were highly stable, and the results did not change with downsampling. Moreover, the percentage of total variance explained by the first PCs remains above 60% for all conditions, which further indicates that the results of the PCA are reliable. The rotation of neural tuning toward the vertical effort axis indicates that the encoding of reward is ‘compressed’ more than is the encoding of effort. Furthermore, the explained variance by the first principal component is significantly decreased, which further indicates an overall compression in utility signaling at higher doses of AMPH, as opposed to saline or low-dose AMPH (Fig.5; ANOVA, F_3,60_ = 17.3454; p = 3 × 10^−8^, bootstrapped 100 times). This effect might be through a breakdown in the encoding of effort and reward or an increased dispersion in effort-reward coding by shifting some neurons to the first quadrant in which the neurons respond positively to both effort and reward. As a result, these neurons fail to encode utility.

**Figure 4.**
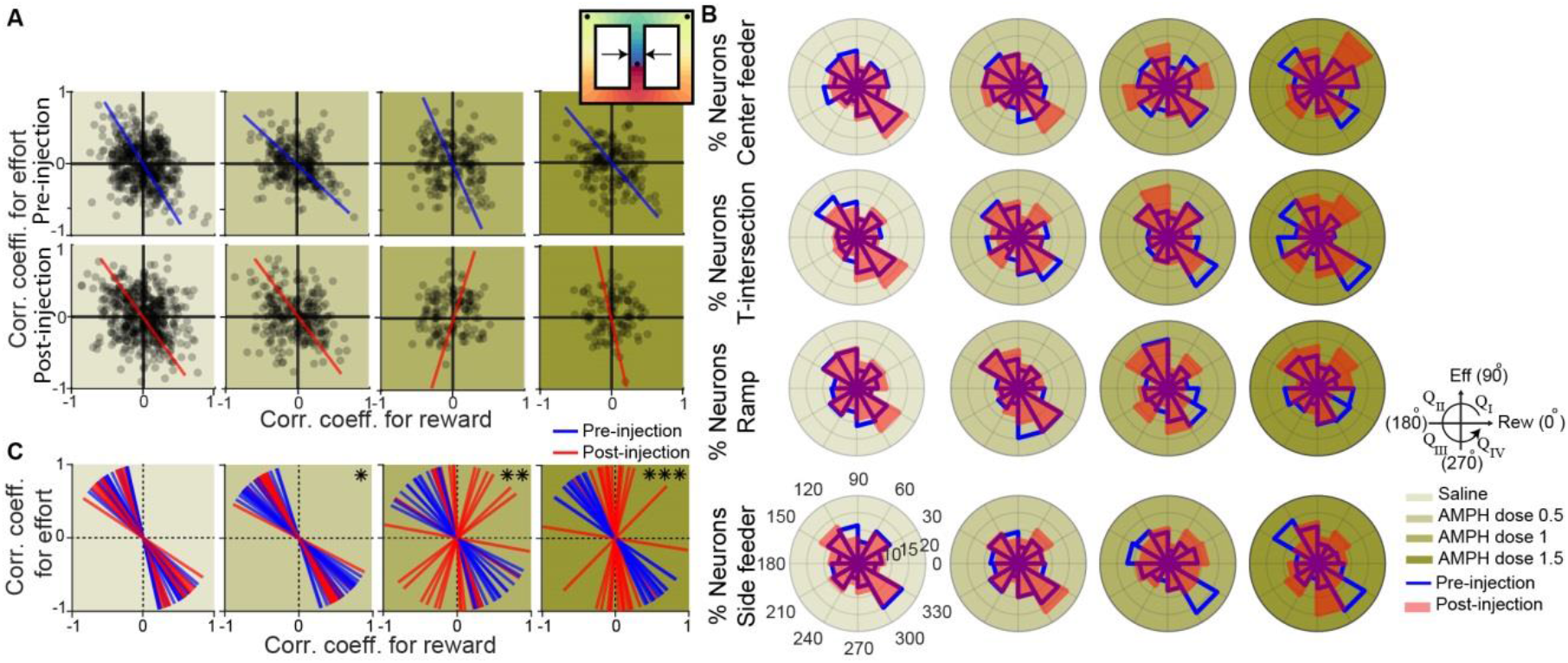
Effect of AMPH on the joint encoding of reward and effort by single neurons. A, Representative example of effort-reward joint encoding at spatial bin #3 (bin adjacent to the central feeder, as indicated by arrows in the inset). Each black circle shows the Pearson correlation coefficient between the amount of effort and firing rate of one neuron, plotted against the correlation value for reward and firing rate of the same neuron. Neurons from all sessions are aggregated. The blue and red lines indicate the first principal component (PC) coefficient of pre- (blue) and post- (red) injection for all units in the dataset. B, Polar distribution of neurons in the effort-reward joint-coding plane for four spatial bins: Central feeder, T-intersection, Barrier, and Side feeder. Angles (0° to 360°) and quadrants (Q_I_ to Q_IV_) are defined as in the plane geometry. The order of quadrants and angles are shown at the right subpanel. C, First PC coefficients of all cells obtained in the 16 spatial bins, from central feeder to a bin after the side feeder for saline or AMPH. (Kuiper two-sample test for pre- vs post-injection, *: significant at p<0.05, **: p<0.01, ***: p <0.001).

**Figure 5.**
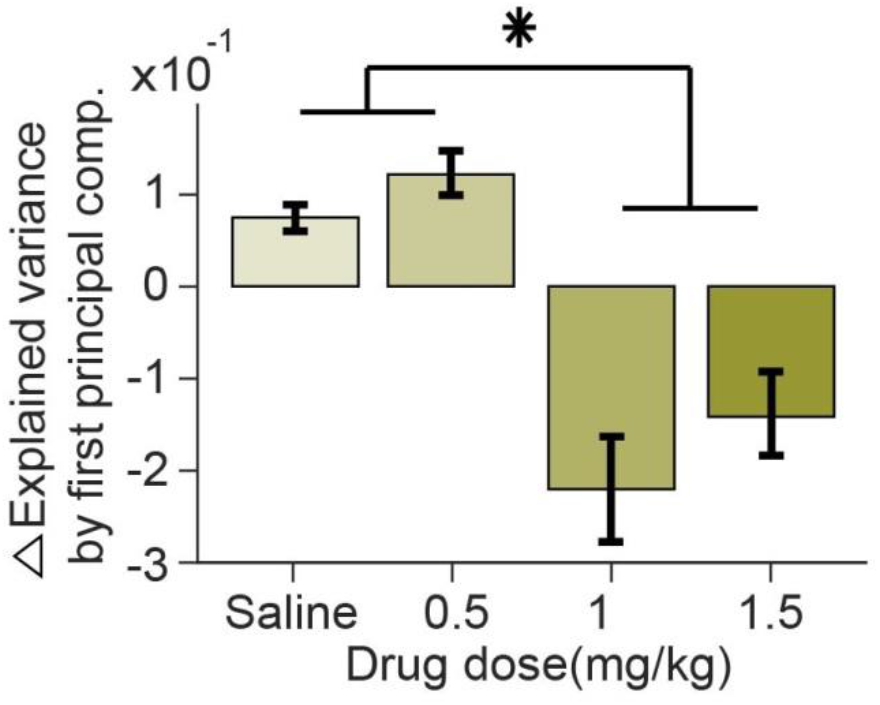
Dose-dependent effect of AMPH on the explained variance of effort-reward encoding. The mean relative change of explained variance of data by the first PC of effort-reward jointly encoding neurons, as well as its SEM. (ANOVA, F_3,60_ = 17.5434; p = 3 × 10^−8^, *: significant at p<0.005 - pairwise comparisons).

In sum, high doses of AMPH induce a significant impairment in the encoding of utility by single units. This effect appears to be caused predominantly by a reduction in reward signaling, rather than effort signaling, which is consistent with our behavioural observations that AMPH-treated rats are less interested in consuming the reward. These results therefore suggest that reward is devalued by AMPH to a greater extent than is effort.

### AMPH contracts ensemble state space

The analysis above suggests that the encoding of utility by single units is compressed by AMPH. This presumes that the primary carrier of information is the mean firing rate of cells, and does not take into account coordination of firing among neurons. We therefore conducted a state-space analysis of simultaneously recorded neurons to assess how AMPH affects information encoded by temporally-evolving patterns of neural ensembles. We used a method termed Gaussian Process Factor Analysis (GPFA) to reduce the dimensionality of the data. This algorithm is particularly advantageous for producing smooth trajectories in low dimensional space from processes with discrete events, such as action potentials space from processes with discrete events, such as action potentials (Yu *et al*., 2009). Neural trajectories obtained by GPFA are modulated by multiple features of the task, including position, effort, and reward. In the 3D space of the first three GPFA factors, neural trajectories are similar for similar trial types, such as those with the same conditions of effort and reward. Figure 6 shows an example of these trajectories for trials of low/high effort in two sessions, and are color coded based on the animal’s location in the maze (Fig.6 inset). The return path, from side feeder to the central feeder, is removed from the Figure to facilitate visualization. Trajectories therefore start at the central feeder (blue) and end after reward consumption at a side feeder (orange). Trajectories deviate significantly among low- and high-effort trials (solid vs dashed lines), particularly while the rat is approaching and climbing the barrier. This shows that the motoric demands of the task modulate the ensemble activity of the neural population, as expected from the single neuron analysis (Fig.3). This indicates that the encoding is specific to task elements other than spatial location. This is further evident by the large deviations in ensemble encoding when animals spontaneously engage in off-task behavior during trials of the task (Fig.7).

**Figure 6.**
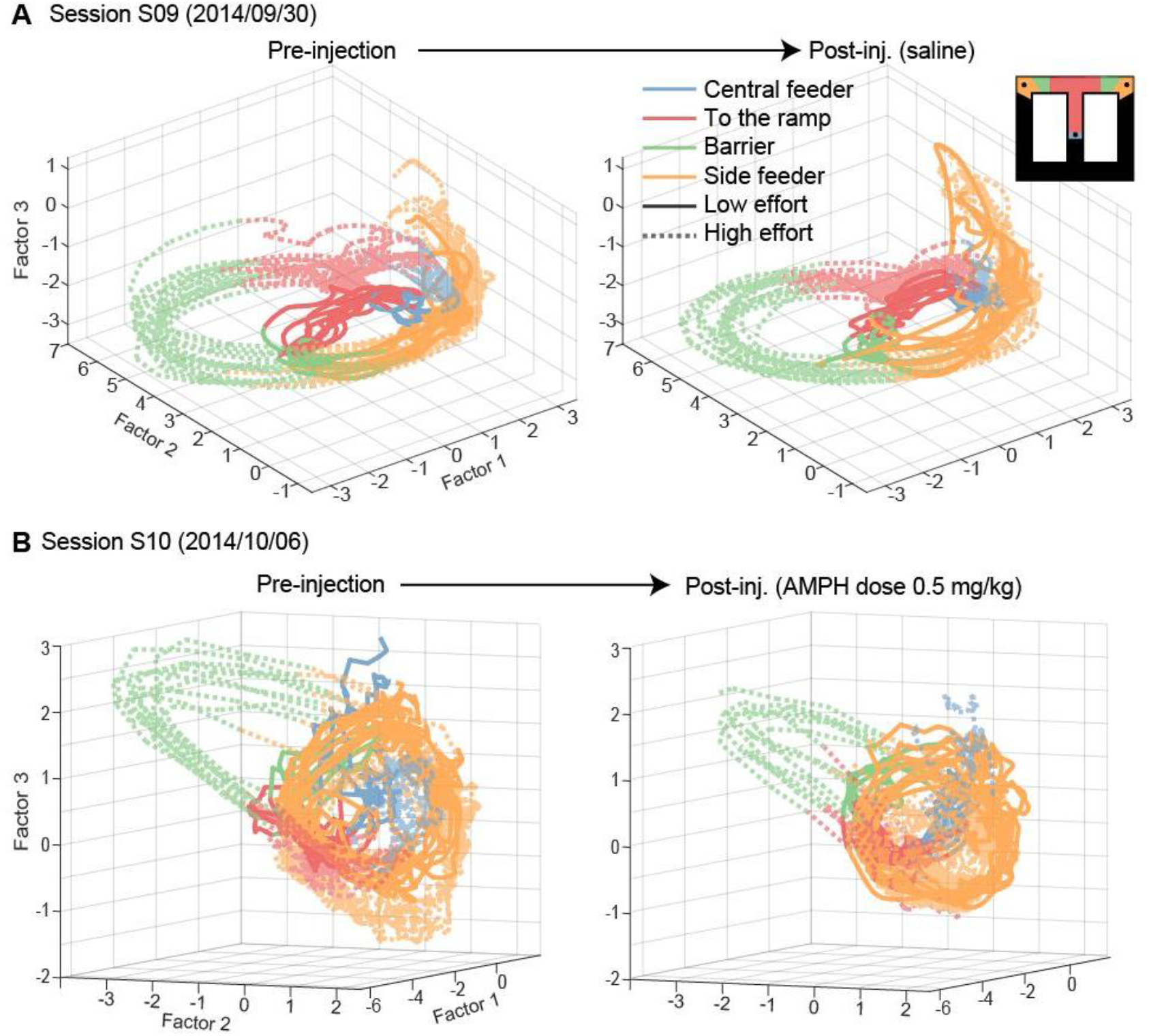
Representative example of effort discrimination via neural trajectories. Neural trajectories of effort trials from the central to side feeders are shown in a reduced 3D state-space before and after the injection of saline (panel A) or AMPH (panel B). The trajectories in A and B are derived from two different sessions in the same animal, and thus show different state-spaces because the population of recorded neurons are distinct. The state-spaces are identical for right and left plots within each panel. The figure inset shows the color-map of the task epochs. High-effort trials are shaded with transparent colors and dotted lines for better visualization.

**Figure 7.**
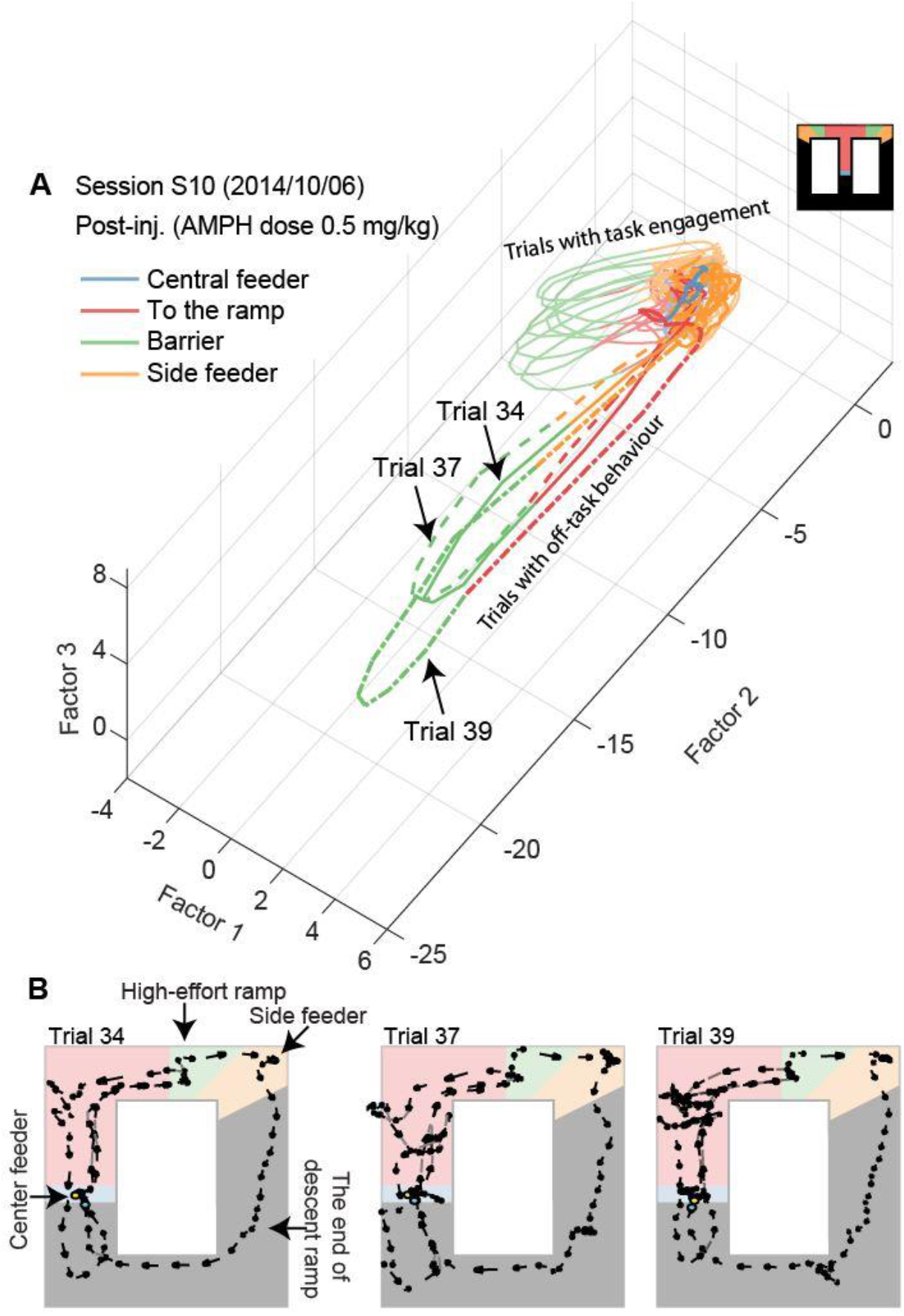
Ensemble encoding deviates largely when rats perform off-task behavior. A, trajectories of population activity in latent space. The trajectory forms overlapping loops in the upper right-hand corner when the animal is performing the task, as in figure 6B. The trajectory deviates widely when the animal wonders on the track, as seen in three exemplars indicated by trial numbers. B, running path of the rat on the trials in panel A with off-task locomotion. The animal approaches the barrier, and then back-tracks toward the starting feeder (blue region) rather than climb the barrier to reach the side feeder (orange region).

Injection of saline had little effect on the trajectories in the reduced-dimension state-space (Fig.6A), whereas AMPH appears to cause a modest contraction in the space (Fig.6B). To better visualize the encoding of task features and the effects of AMPH, we next independently plotted the first four GPFA factors to compare encoding on trials with high-vs low-reward (and constant effort; Fig.8). The first factor in each session shows a clear discrimination of reward at the feeder zones. In some cases, the second factor does as well. AMPH does not appear to strongly affect the discrimination by the ensemble, but does appear to reduce the overall variance of factors (peak to peak amplitude). Note that the roughness of some trajectories (e.g. Fig.8B and C as compared to A and D) might be due to the lower number of cells in those sessions or increased population correlation and is not the effect of drug conditions as they appear even before the injections.

**Figure 8.**
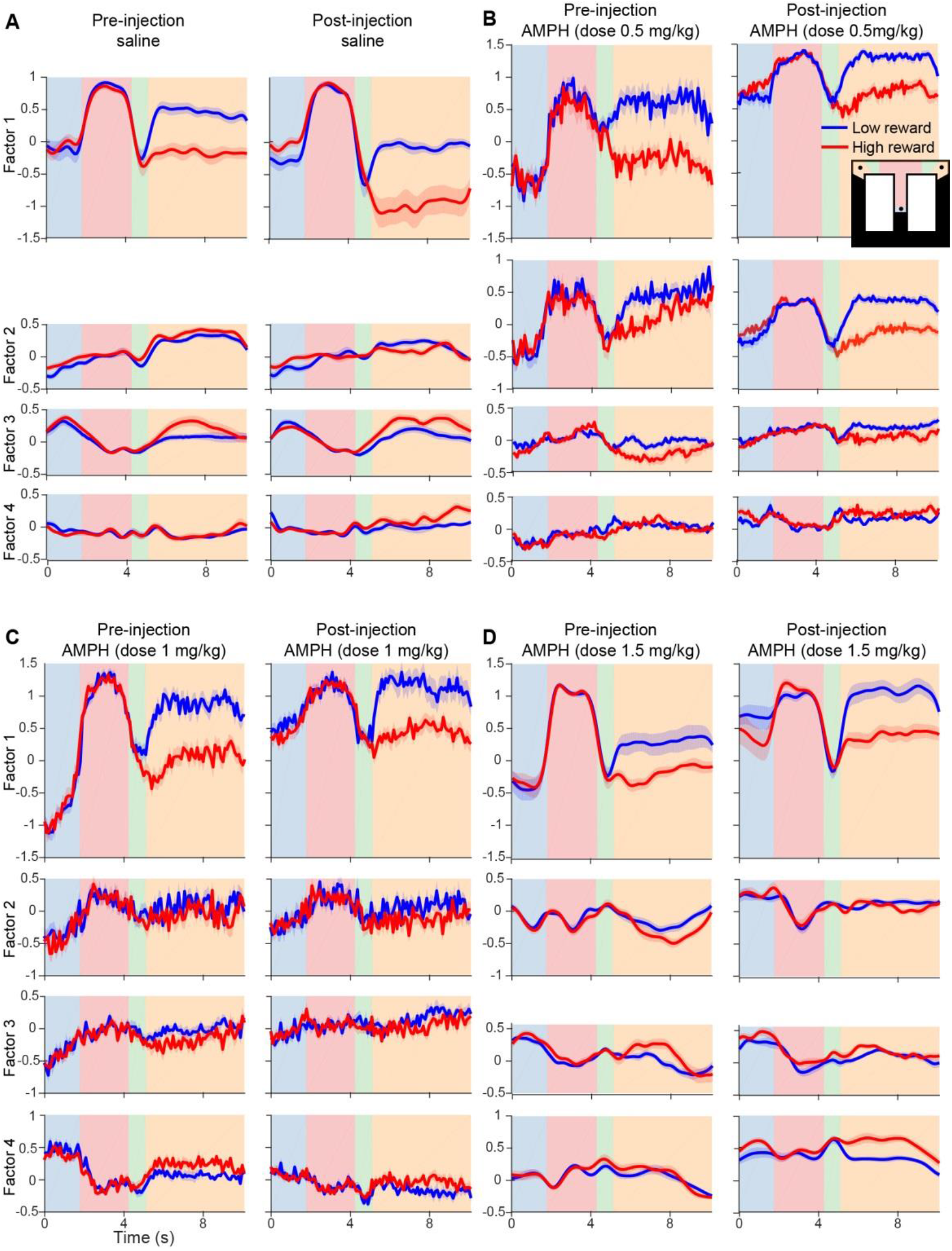
Representative example of reward discrimination by ensemble activity. A, The first four latent factors of population activity in low- and high-reward trials (color) are shown as the rat traverses from central to side feeders (linearized time in horizontal axis) before and after injection of saline. The solid lines indicate the mean, and the shaded area represents the SEM. Background colours indicate the task epoch as shown in the inset of panel B. The duration of each epoch is normalized prior to producing group statistics. B, C, and D, show data from the same rat for AMPH doses of 0.5, 1, and 1.5 mg/kg, respectively.

We next sought to quantitatively test these observations. Using only data from bins between the central feeder to target feeders, we tested if the level of effort or reward could be discriminated by the mean value of either the first or second GPFA factor across each task epoch (ANOVA, significant at p<0.05). We then computed the fraction of sessions with significant effort or reward encoding. The effort is well discriminated at all epochs (Fig.9A) in most sessions (>50%), but peaks significantly near the barrier (ANOVA, F_3,28_ = 6.6; p = 0.0016 for first, and F_3,28_ = 19.01; p = 6×10^−7^ for second factor). The reward, on the other hand, is only discriminated well at the target feeders (Fig.9B) (ANOVA, F_3,28_ = 47.55; p = 4× 10^−11^ for first, and F_3,28_ = 31.41; p = 4× 10^−9^ for second factor). AMPH does not affect the discrimination of effort or reward (Fisher’s exact test, p>0.05 for pre-vs post-injection of saline or drug in each epoch). It is interesting to note that although the relative fraction of cells encoding reward is low and does not change across task epochs (Fig.3), the information encoded by the population is strongly modulated by task epochs, and discriminates reward amount with high accuracy (>80%) at the side feeder.

**Figure 9.**
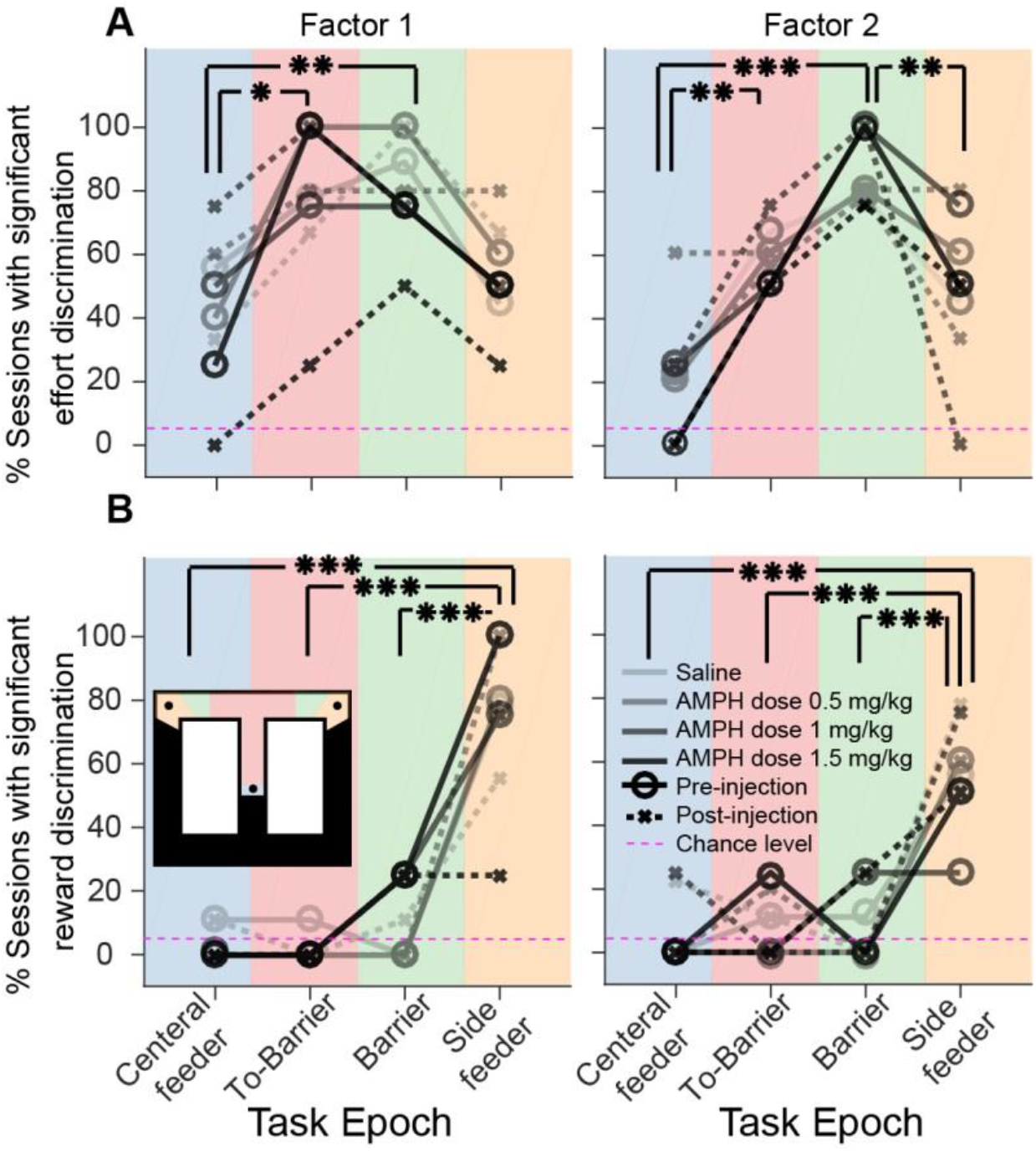
Effort and reward encoding by neural ensembles. Mean proportion of sessions in which the conditional mean of the first or second GPFA factors can discriminate effort (panel A) or reward (panel B) before or after injection. Each drug or saline condition is distinguished by a specific line opacity. Solid lines and circular markers indicate the percentage of sessions in which the given parameter (effort or reward) significantly modulated pre-injection GPFA factors; dotted lines of the same shade with ‘x’ markers indicate the same variable for the post-injection phase of the same condition. The background color indicates the four task epochs analyzed as shown in the inset: central feeder, approach to the barrier, climbing the barrier, and side feeder. (ANOVA, *: significant at p<0.01, **: p<0.005, ***: p <0.0001).

To test if AMPH contracted the trajectories in state-space, we computed the change by AMPH of the volume occupied by the GPFA factor trajectories in high- and low-reward trials projected into the same state-space (Fig.10). As compared to saline, the volume is significantly reduced by 0.5 and 1.0 mg/kg of AMPH in low-reward condition (RM-ANOVA, F_2.465,93.68_ = 12.267, p = 4×10^−6^) and by 0.5 and 1.0 mg/kg in high-reward condition (RM-ANOVA, F_1.729,67.437_ = 10.073, p = 3 × 10^−4^). It is possible that the increased variance in the rat’s movement path and/or increased propensity for off-task behaviour in the high-dose condition may obscure the direct pharmacological effects in the ACC by increasing unexplained variance.

**Figure 10.**
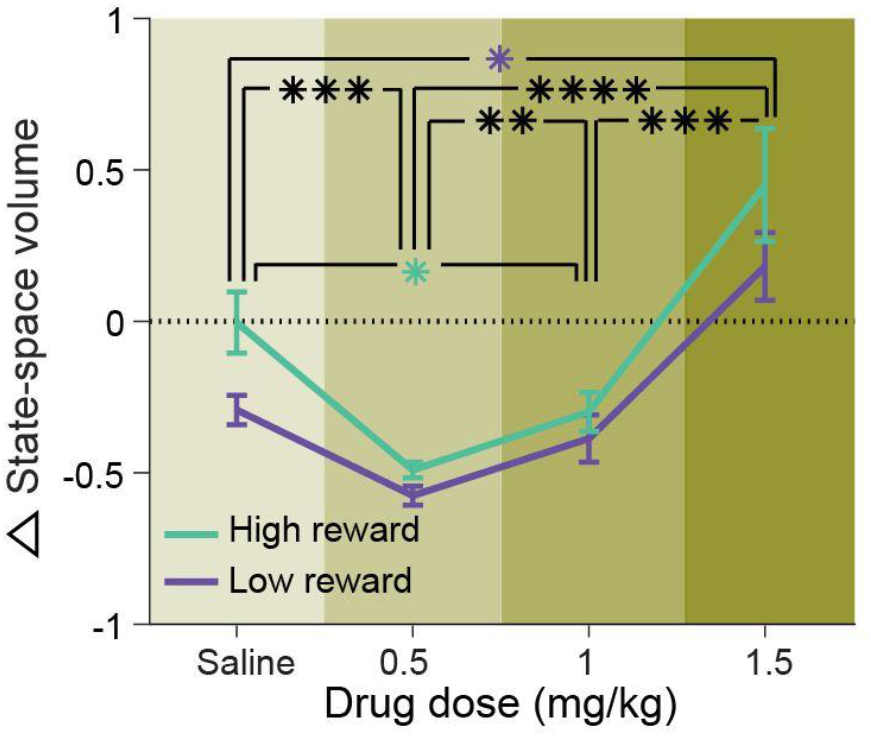
Dose-dependent effect of AMPH on state-space volume. The mean relative volume changes in high- and low-reward trials, as well as its SEM. The volume is calculated as the state-space enclosed by the neural trajectory from central to side feeders in the 3D space formed by the first three latent factors of the pre-injection condition. The volumes of all post-injection trials were then compared with the mean volume of the same group of trials in pre-injection phase. This comparison occurs in the same space, which is specific to the session, and then the relative change is compared across the sessions as shown here. Negative and positive values in relative change demonstrate contraction and expansion in the state-space volume, respectively. (RM-ANOVA, *: significant at p<0.05, **: p<0.01, ***: p <0.005 and ****: p<10^−6^)

We lastly analyzed the dynamics of the ensemble by computing the state-space decorrelation, which describes how the correlation of vectors of binned neural activity change in time and space (McNaughton, 1998; Battaglia *et al*., 2004). The state vector correlations are shifted upward, indicating that the patterns are more correlated across the entire maze (Fig. 11A). Only the 1.5 mg/kg dose showed a statistically significant difference from the saline condition (ANOVA, F_3,18_ = 6.97; p = 0.0026). For this high dose, the half-amplitude of the decorrelation is higher, meaning that the rate of change of correlation is lower (Fig. 11B).

**Figure 11.**
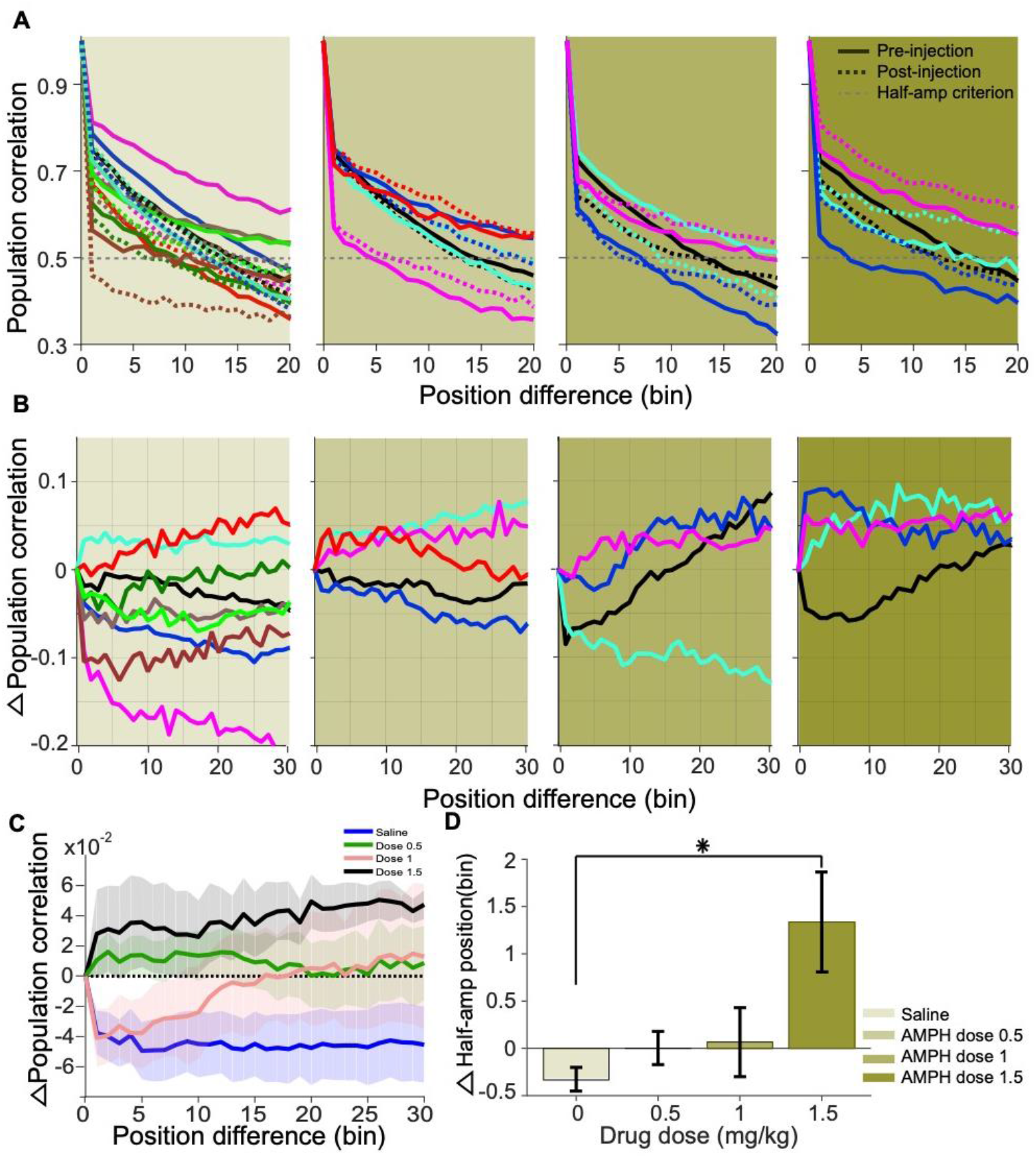
Effect of AMPH on population vector decorrelation. A, Population vector correlation of all task epochs of pre- and post-injection for each session (distinguished by color; such that each solid line indicates pre-injection population correlation and dotted line of the same color indicates the post-injection correlation function of the same session). Grey dotted horizontal lines represent the half-amplitude criteria. The crossing of this line indicates how fast the population decorrelates. B, Differences of population vector correlation after injection with respect to before, which corresponds to the differences of dotted lines from solid lines of the panels in A. C, Mean change in population vector correlation (mean of lines in panel B), as well as their SEM. D, Mean relative change in the half-amplitude crossing point following injections. (Errorbars: SEM and *: significant at p<0.005, ANOVA).

In sum, the first and second latent GPFA factors indicate that the neural ensemble is strongly modulated prior to the effort (approaching to the barrier), while exerting effort (climbing the barrier), and while at the feeder. Although AMPH did not affect the mean difference of the first two GPFA factors based on effort or reward, intermediate doses did decrease the state space volume.

## Discussion

In this study, we investigated the effect of AMPH on ACC neural activity in rats performing a task with variable effort and reward. Our results show that individual neurons jointly encoding effort and reward are affected by AMPH. This joint encoding is expected of cells involved in computing utility. The activity of most units is positively correlated with the reward amount and negatively with the effort level during all task epochs, form the central start feeder to the side feeders. AMPH injection increased the dispersion of effort-reward coding in the population, indicating some deterioration of utility encoding in terms of both reward and effort. Nonetheless, the rotation of the principle axis of the population toward the effort axis by AMPH reveals that the suppression of reward signaling by single neurons is the primary effect.

The analysis of ensemble activity revealed a clear event-specific discrimination of effort and reward levels. Consistent with the single-unit analysis, the effort discrimination was maximal in the proximity of the barrier, whereas reward discrimination was largest at the side feeders. Although the discrimination of effort and reward at the population level was not attenuated by AMPH, the amplitude of signaling within the latent space (after dimensionality reduction) was reduced by intermediate doses of AMPH, leading to a reduced volume of neural trajectories in the latent space. Moreover, the spatial decorrelation of neural patterns was reduced by increasing doses of AMPH. These data reveal that AMPH causes a contraction of signaling and an increase in self-similarity over longer distances. This is consistent with the analyses of single units, which indicate that the contraction may be more pronounced in the reward domain. A loss of reward-related variance without increase in variance related to other task features will manifest as increased self-similarity because reward signaling is not uniform across epochs.

### AMPH and the perception of choice value

How might the above-described effects of AMPH on neural signaling affect ACC function? The systemic injection of AMPH has widespread effects in the brain. Catecholamines in several brain regions have been postulated to influence action generation and reward learning (Berridge, 2007), and effortful responses for goal-directed behaviours (Salamone & Correa, 2002). Part of the effect may be locomotor. AMPH increases motoric activity (Randrup & Munkvad, 1967; Groves & Rebec, 1976; Robinson & Berridge, 1993; Wilkinson *et al*., 1993; Sams-Dodd, 1998), an effect we observed here. Moreover, AMPH has been shown to decrease feeding (Foltin, 2001; Shoblock *et al*., 2003; Cannon *et al*., 2004; Wellman *et al*., 2009) and sensitivity to reward omission (Wong *et al*., 2017). Consistent with these data, we found that animals spent less time at feeders, and were more likely to forgo the reward, as AMPH increased. Thus, the behavioural effects appear dichotomous; animals increased task engagement, but were less interested in the reward. This pattern of behavioural effects may reflect discounting of both the physical effort and reward value, but then it remains unclear why an animal would work for a highly discounted outcome, rather than engage in some other behaviour. Indeed, animals did not engage in the task at higher doses (≥ 2.0 mg/kg), and instead engaged in off-task behaviours such as grooming, sniffing, and exploring the edges of the track. A similar pattern of behavioural effects have been reported elsewhere. For instance, moderate AMPH significantly improves working memory and increases locomotor activity, whereas higher doses do not (Shoblock *et al*., 2003). We later speculate why AMPH has this dose dependency and seemingly paradoxical effect on reward value.

We cannot infer from our data where in the brain AMPH may be affecting decision-related processing. We can, however, use neural activity in ACC as a window into network processing. Previous studies have reported ACC activity in anticipation of effort and reward (Sul *et al*., 2010; Cowen *et al*., 2012; Hashemniayetorshizi *et al*., 2015). Our results show that most neurons primarily encode utility in the portion of the maze from the center feeder to side feeders. It is possible that the animals anticipate the effort and reward of the upcoming trial because of the task design; the animal was directed to alternate between left and right options, and the effort and rewards were static over blocks of 20 trials. Animals are thus not able to choose among the options, so the ‘choice’ is between performing and not performing the task. It is therefore possible that modulation of neural activity reported here serves to track expectations or keep the animal engaged in the task, rather than generating optimal cost-benefit decisions among the right and left feeders. AMPH may either change the value and/or neural mechanisms of task engagement, or could affect the mechanisms by which the animal maintains engagement. This function is more akin to attention and vigilance, which are both increased by AMPH in a variety of task settings and species (Sostek *et al*., 1980; Ridley *et al*., 1982; Koelega, 1993; Solanto, 1998; Grilly, 2000; Silber *et al*., 2006; Sagvolden, 2011). These phenomena are likely a property of network dynamics rather than signal encoding by individual units, as discussed next.

### AMPH contracts populations neural dynamics

ACC population activity has been reported to evolve through a stereotyped set of states during task performance such that each task epoch is encoded by a specific encoding state (Lapish *et al*., 2008; Balaguer-Ballester *et al*., 2011). We likewise found that the ensembles followed a stereotyped trajectory in latent space, and that the trajectory deviated widely when rats were not performing the task (Fig.7). These data suggest that the ACC encoding evolves through a neural ‘script’ when on-task. Past computational models have suggested that dopamine stabilizes the dynamics in the prefrontal cortex by direct neuromodulatory effects on cortical neurons (Gruber *et al*., 2006; Durstewitz & Seamans, 2008), as well as its afferents (Gruber *et al*., 2006). This provides an explanatory framework by which AMPH could produce the observed effects in our task. Neuromodulation by low-dose AMPH stabilizes task-related patterns, which is predicted by the models to decrease the probability of task disengagement. Note that this mechanism requires no notion of reward value or utility; rather, it is a property of changing the neural dynamics in the network. Furthermore, AMPH has been reported to increase the separation between distinct ensemble activity patterns during distinct task states at a moderate dose (1 mg/kg), whereas it reduces the distance between such distinct neural activity states at high doses (3.3 mg/kg) (Lapish *et al*., 2015). Our results are consistent with this, but with the added feature that the occupancy of trajectories in state space is also affected. At low doses (0.5-1.0 mg/kg), we found a slight contraction of ensemble state space. This became an expansion of state-space at high doses (1.5 mg/kg). Moreover, the decorrelation time increased at high doses. These observations are consistent with that of Lapish and colleagues. Although the state space contracts under low doses, the variance from trial to trial also appears to reduce, which can increase pattern separation from samples at different points of the trajectory (different task epochs). At high doses, the state space expands, but so does the variance across trials. The increased variance and increased correlation across location both increase the similarity (overlap) of pattern samples at different task epochs. In other words, the decrease in uninformative variance (i.e. ‘noise’) under low-dose AMPH is proportionally greater than the decrease in state space contraction of ensemble patterns across phases of the task, thus leading to a greater separation of patterns of distinct task epochs (in the sense of discrimination analyses, such as D prime in signal detection theory). This breaks down under higher doses of AMPH such that the encoding becomes plagued with variance that can knock the state of ACC encoding off of its task-related trajectory, thus deviating from the behavioral script and breaking engagement in the task. It may be that at high doses, only very robust behaviours can withstand the additional variance, such that the behavioural output consists of a repertoire of innate behaviours and ‘over-learned’ operant responses. Other less stable responses may begin to be initiated, but are disrupted by the high variance. Indeed, the behavioural output under high doses of AMPH and other psychostimulants are typically highly stereotyped actions reflecting parts of grooming, sniffing, and licking with repetitive head movement (Randrup & Munkvad, 1967; Fog, 1970; Schiorring, 1971). Our interpretation presumes that neural activity in ACC reflects neural dynamics elsewhere in the brain that produce behavior. We cannot rule out the inverse relationship – that some other brain system drives the behavior, and the ACC encoding tracks the animal’s state. Our data are consistent with past results showing that rat ACC activity is sensitive to small deviations of position on a track (Euston & McNaughton, 2006). It is therefore possible that AMPH acts on a brain system uncorrelated with ACC dynamics to produce increased variance in the running path, which then increases variance in ACC dynamics. Our data provide some evidence against this possibility. AMPH has a monotonically increasing effect on running path roughness, but reduces ACC variance at low-to-moderate doses. This suggests that ACC encoding is not dictated by the positional state of the animal.

Although our data are purely correlational, they suggest novel linkages between the effects of AMPH on neural activity and behaviour. We propose that the reason rats perform the task even though they lose interest in the reward under moderate doses of AMPH is because of this drug’s combined effects on reward encoding and dynamics. Specifically, rats have less interest in the reward because reward information is attenuated. Nonetheless, rats continue to engage in the task at moderate AMPH levels because the ensemble dynamics are stabilized such that rats are less likely to disengage. The result is that the brain runs quickly through the task script, even though the reward is not desired.

## Acknowledgements

This work was supported by the National Sciences and Engineering Research Council of Canada (NSERC) and Alberta Innovates Health Solutions.

## Ethics

Animal experimentation: All procedures were approved by the university’s animal welfare committee (Protocol # 1502) in accordance with the Canadian Council on Animal Care.

## References

Amemori, K. & Graybiel, A.M. (2012) Localized microstimulation of primate pregenual cingulate cortex induces negative decision-making. Nat Neurosci, 15, 776–785.

Balaguer-Ballester, E., Lapish, C.C., Seamans, J.K. & Durstewitz, D. (2011) Attracting dynamics of frontal cortex ensembles during memory-guided decision-making. PLoS Comput Biol, 7, e1002057.

Bardgett, M.E., Depenbrock, M., Downs, N., Points, M. & Green, L. (2009) Dopamine modulates effort-based decision making in rats. Behav Neurosci, 123, 242–251.

Bartho, P., Hirase, H., Monconduit, L., Zugaro, M., Harris, K.D. & Buzsaki, G. (2004) Characterization of neocortical principal cells and Interneurons by network interactions and extracellular features. J Neurophysiol, 92, 600–608.

Battaglia, F.P., Sutherland, G.R. & McNaughton, B.L. (2004) Local sensory cues and place cell directionality: Additional evidence of prospective coding in the hippocampus. Journal of Neuroscience, 24, 4541–4550.

Berridge, K.C. (2007) The debate over dopamine’s role in reward: the case for incentive salience. Psychopharmacology, 191, 391–431.

Blanchard, T.C. & Hayden, B.Y. (2014) Neurons in dorsal anterior cingulate cortex signal postdecisional variables in a foraging task. J Neurosci, 34, 646–655.

Blanchard, T.C., Strait, C.E. & Hayden, B.Y. (2015) Ramping ensemble activity in dorsal anterior cingulate neurons during persistent commitment to a decision. J Neurophysiol, 114, 2439–2449.

Blundell, J.E., Latham, C.J. & Leshem, M.B. (1976) Differences between Anorexic Actions of Amphetamine and Fenfluramine-Possible Effects on Hunger and Satiety. J Pharm Pharmacol, 28, 471–477.

Blundell, J.E., Latham, C.J., Moniz, E., Mcarthur, R.A. & Rogers, P.J. (1979) Structural-Analysis of the Actions of Amphetamine and Fenfluramine on Food-Intake and Feeding-Behavior in Animals and in Man. Curr Med Res Opin, 6, 34–54.

Cannon, C.M., Abdallah, L., Tecott, L.H., During, M.J. & Palmiter, R.D. (2004) Dysregulation of striatal dopamine signaling by amphetamine inhibits feeding by hungry mice. Neuron, 44, 509–520.

Chiueh, C.C. & Moore, K.E. (1973) Release of Endogenously Synthesized Catechols from Caudate-Nucleus by Stimulation of Nigro-Striatal Pathway and by Administration of D-Amphetamine. Brain Research, 50, 221–225.

Cousins, M.S., Atherton, A., Turner, L. & Salamone, J.D. (1996) Nucleus accumbens dopamine depletions alter relative response allocation in a T-maze cost/benefit task. Behav Brain Res, 74, 189–197.

Cowen, S.L., Davis, G.A. & Nitz, D.A. (2012) Anterior cingulate neurons in the rat map anticipated effort and reward to their associated action sequences. J Neurophysiol, 107, 2393–2407.

Cowen, S.L. & McNaughton, B.L. (2007) Selective delay activity in the medial prefrontal cortex of the rat: contribution of sensorimotor information and contingency. J Neurophysiol, 98, 303–316.

Croxson, P.L., Walton, M.E., O’Reilly, J.X., Behrens, T.E.J. & Rushworth, M.F.S. (2009) Effort-Based Cost-Benefit Valuation and the Human Brain. Journal of Neuroscience, 29, 4531–4541.

Durstewitz, D. & Seamans, J.K. (2008) The Dual-State Theory of Prefrontal Cortex Dopamine Function with Relevance to Catechol-O-Methyltransferase Genotypes and Schizophrenia. Biol Psychiat, 64, 739–749.

Durstewitz, D., Vittoz, N.M., Floresco, S.B. & Seamans, J.K. (2010) Abrupt transitions between prefrontal neural ensemble states accompany behavioral transitions during rule learning. Neuron, 66, 438–448.

Euston, D.R. & McNaughton, B.L. (2006) Apparent encoding of sequential context in rat medial prefrontal cortex is accounted for by behavioral variability. J Neurosci, 26, 13143–13155.

Floresco, S.B., Tse, M.T. & Ghods-Sharifi, S. (2008) Dopaminergic and glutamatergic regulation of effort- and delay-based decision making. Neuropsychopharmacology, 33, 1966–1979.

Fog, R. (1970) Behavioural effects in rats of morphine and amphetamine and of a combination of the two drugs. Psychopharmacologia, 16, 305–312.

Foltin, R.W. (2001) Effects of amphetamine, dexfenfluramine, diazepam, and other pharmacological and dietary manipulations on food “seeking” and “taking” behavior in non-human primates. Psychopharmacology, 158, 28–38.

Fujisawa, S., Amarasingham, A., Harrison, M.T. & Buzsaki, G. (2008) Behavior-dependent short-term assembly dynamics in the medial prefrontal cortex. Nat Neurosci, 11, 823–833.

Gneiting, T., Sevcikova, H. & Percival, D.B. (2012) Estimators of Fractal Dimension: Assessing the Roughness of Time Series and Spatial Data. Stat Sci, 27, 247–277.

Grilly, D.M. (2000) A verification of psychostimulant-induced improvement in sustained attention in rats: Effects of d-amphetamine, nicotine, and pemoline. Exp Clin Psychopharm, 8, 14–21.

Groves, P.M. & Rebec, G.V. (1976) Biochemistry and behavior: some central actions of amphetamine and antipsychotic drugs. Annu Rev Psychol, 27, 91–127.

Gruber, A.J., Calhoon, G.G., Shusterman, I., Schoenbaum, G., Roesch, M.R. & O’Donnell, P. (2010) More is less: a disinhibited prefrontal cortex impairs cognitive flexibility. J Neurosci, 30, 17102–17110.

Gruber, A.J., Dayan, P., Gutkin, B.S. & Solla, S.A. (2006) Dopamine modulation in the basal ganglia locks the gate to working memory. J Comput Neurosci, 20, 153–166.

Gruber, A.J., Hussain, R.J. & O’Donnell, P. (2009) The nucleus accumbens: a switchboard for goal-directed behaviors. PLoS One, 4, e5062.

Hashemniayetorshizi, S., Sessford, D., Gruber, A.J. & Euston, D.R. (2015) Modulation of cellular activity by effort and reward in Anterior Cingulate Cortex. Program No. 633.20. 2015 Neuroscience Meeting Planner. Washington, DC: Society for Neuroscience, *2015. Online*..

Hauber, W. & Sommer, S. (2009) Prefrontostriatal circuitry regulates effort-related decision making. Cereb Cortex, 19, 2240–2247.

Hausdorff, F. (1919) Dimension und äußeres Maß. Mathematische Annalen, 79, 157–179.

Hayden, B.Y., Pearson, J.M. & Platt, M.L. (2009) Fictive Reward Signals in the Anterior Cingulate Cortex. Science, 324, 948–950.

Hillman, K.L. & Bilkey, D.K. (2010) Neurons in the rat anterior cingulate cortex dynamically encode cost-benefit in a spatial decision-making task. J Neurosci, 30, 7705–7713.

Holec, V., Pirot, H.L. & Euston, D.R. (2014) Not all effort is equal: the role of the anterior cingulate cortex in different forms of effort-reward decisions. Front Behav Neurosci, 8, 12.

Isomura, Y., Ito, Y., Akazawa, T., Nambu, A. & Takada, M. (2003) Neural coding of “attention for action” and “response selection” in primate anterior cingulate cortex. J Neurosci, 23, 8002–8012.

Kennerley, S.W., Dahmubed, A.F., Lara, A.H. & Wallis, J.D. (2009) Neurons in the frontal lobe encode the value of multiple decision variables. J Cogn Neurosci, 21, 1162–1178.

Kennerley, S.W. & Wallis, J.D. (2009) Evaluating choices by single neurons in the frontal lobe: outcome value encoded across multiple decision variables. Eur J Neurosci, 29, 2061–2073.

Kennerley, S.W., Walton, M.E., Behrens, T.E., Buckley, M.J. & Rushworth, M.F. (2006) Optimal decision making and the anterior cingulate cortex. Nat Neurosci, 9, 940–947.

Klein-Flugge, M.C., Kennerley, S.W., Friston, K. & Sestmann, S. (2016) Neural Signatures of Value Comparison in Human Cingulate Cortex during Decisions Requiring an Effort-Reward Trade-off. Journal of Neuroscience, 36, 10002–10015.

Koelega, H.S. (1993) Stimulant-Drugs and Vigilance Performance - a Review. Psychopharmacology, 111, 1–16.

Lapish, C.C., Balaguer-Ballester, E., Seamans, J.K., Phillips, A.G. & Durstewitz, D. (2015) Amphetamine Exerts Dose-Dependent Changes in Prefrontal Cortex Attractor Dynamics during Working Memory. J Neurosci, 35, 10172–10187.

Lapish, C.C., Durstewitz, D., Chandler, L.J. & Seamans, J.K. (2008) Successful choice behavior is associated with distinct and coherent network states in anterior cingulate cortex. Proc Natl Acad Sci U S A, 105, 11963–11968.

Leibowitz, S.F., Shorposner, G., Maclow, C. & Grinker, J.A. (1986) Amphetamine - Effects on Meal Patterns and Macronutrient Selection. Brain Res Bull, 17, 681–689.

Mai, B., Sommer, S. & Hauber, W. (2012) Motivational states influence effort-based decision making in rats: the role of dopamine in the nucleus accumbens. Cogn Affect Behav Neurosci, 12, 74–84.

Mashhoori, A., Hashemnia, S., McNaughton, B.L., Euston, D.R. & Gruber, A.J. (2018) Rat anterior cingulate cortex recalls features of remote reward locations after disfavoured reinforcements. Elife, 7.

McNaughton, B.L. (1998) The neurophysiology of reminiscence. Neurobiol Learn Mem, 70, 252–267.

McNaughton, B.L., O’Keefe, J. & Barnes, C.A. (1983) The stereotrode: a new technique for simultaneous isolation of several single units in the central nervous system from multiple unit records. J Neurosci Methods, 8, 391–397.

Odum, A.L. & Shahan, T.A. (2004) D-Amphetamine reinstates behavior previously maintained by food: importance of context. Behav Pharmacol, 15, 513–516.

Paxinos, G. & Watson, C. (2014) Paxino’s and Watson’s The rat brain in stereotaxic coordinates. Elsevier/AP, Academic Press is an imprint of Elsevier, Amsterdam; Boston.

Phillips, P.E.M., Walton, M.E. & Jhou, T.C. (2007) Calculating utility: preclinical evidence for cost-benefit analysis by mesolimbic dopamine. Psychopharmacology, 191, 483–495.

Porter, B.S., Hillman, K.L. & Bilkey, D.K. (2019) Anterior cingulate cortex encoding of effortful behavior. J Neurophysiol, 121, 701–714.

Pum, M., Carey, R.J., Huston, J.P. & Muller, C.P. (2007) Dissociating effects of cocaine and d-amphetamine on dopamine and serotonin in the perirhinal, entorhinal, and prefrontal cortex of freely moving rats. Psychopharmacology (Berl), 193, 375–390.

Randrup, A. & Munkvad, I. (1967) Stereotyped activities produced by amphetamine in several animal species and man. Psychopharmacologia, 11, 300–310.

Randrup, A., Munkvad, I. & Udsen, P. (1963) Adrenergic Mechanisms and Amphetamine Induced Abnormal Behaviour. Acta Pharmacol Toxicol (Copenh), 20, 145–157.

Ridley, R.M., Baker, H.F., Owen, F., Cross, A.J. & Crow, T.J. (1982) Behavioral and Biochemical Effects of Chronic Amphetamine Treatment in the Vervet Monkey. Psychopharmacology, 78, 245–251.

Robinson, T.E. & Berridge, K.C. (1993) The Neural Basis of Drug Craving - an Incentive-Sensitization Theory of Addiction. Brain Res Rev, 18, 247–291.

Sagvolden, T. (2011) Impulsiveness, overactivity, and poorer sustained attention improve by chronic treatment with low doses of l-amphetamine in an animal model of Attention-Deficit/Hyperactivity Disorder (ADHD). Behav Brain Funct, 7.

Salamone, J.D. & Correa, M. (2002) Motivational views of reinforcement: implications for understanding the behavioral functions of nucleus accumbens dopamine. Behav Brain Res, 137, 3–25.

Salamone, J.D., Cousins, M.S. & Bucher, S. (1994) Anhedonia or Anergia - Effects of Haloperidol and Nucleus-Accumbens Dopamine Depletion on Instrumental Response Selection in a T-Maze Cost-Benefit Procedure. Behav Brain Res, 65, 221–229.

Sams-Dodd, F. (1998) Effects of continuous D-amphetamine and phencyclidine administration on social behaviour, stereotyped behaviour, and locomotor activity in rats. Neuropsychopharmacology, 19, 18–25.

Schiorring, E. (1971) Amphetamine Induced Selective Stimulation of Certain Behaviour Items with Concurrent Inhibition of Others in an Open-Field Test with Rats. Behaviour, 39, 1-+.

Schweimer, J. & Hauber, W. (2005) Involvement of the rat anterior cingulate cortex in control of instrumental responses guided by reward expectancy. Learn Mem, 12, 334–342.

Schweimer, J. & Hauber, W. (2006) Dopamine D1 receptors in the anterior cingulate cortex regulate effort-based decision making. Learn Mem, 13, 777–782.

Shin, R., Cao, J., Webb, S.M. & Ikemoto, S. (2010) Amphetamine administration into the ventral striatum facilitates behavioral interaction with unconditioned visual signals in rats. PLoS One, 5, e8741.

Shoblock, J.R., Sullivan, E.B., Maisonneuve, I.M. & Glick, S.D. (2003) Neurochemical and behavioral differences between d-methamphetamine and d-amphetamine in rats. Psychopharmacology (Berl), 165, 359–369.

Silber, B.Y., Croft, R.J., Papafotiou, K. & Stough, C. (2006) The acute effects of d-amphetamine and methamphetamine on attention and psychomotor performance. Psychopharmacology, 187, 154–169.

Skvortsova, V., Palminteri, S. & Pessiglione, M. (2014) Learning To Minimize Efforts versus Maximizing Rewards: Computational Principles and Neural Correlates. Journal of Neuroscience, 34, 15621–15630.

Solanto, M.V. (1998) Neuropsychopharmacological mechanisms of stimulant drug action in attention-deficit hyperactivity disorder: a review and integration. Behav Brain Res, 94, 127–152.

Sostek, A.J., Buchsbaum, M.S. & Rapoport, J.L. (1980) Effects of Amphetamine on Vigilance Performance in Normal and Hyperactive-Children. J Abnorm Child Psych, 8, 491–500.

Sul, J.H., Kim, H., Huh, N., Lee, D. & Jung, M.W. (2010) Distinct roles of rodent orbitofrontal and medial prefrontal cortex in decision making. Neuron, 66, 449–460.

Walton, M.E., Bannerman, D.M., Alterescu, K. & Rushworth, M.F.S. (2003) Functional specialization within medial frontal cortex of the anterior cingulate for evaluating effort-related decisions. Journal of Neuroscience, 23, 6475–6479.

Walton, M.E., Bannerman, D.M. & Rushworth, M.F. (2002) The role of rat medial frontal cortex in effort-based decision making. J Neurosci, 22, 10996–11003.

Wellman, P.J., Davis, K.W., Clifford, P.S., Rothman, R.B. & Blough, B.E. (2009) Changes in feeding and locomotion induced by amphetamine analogs in rats. Drug Alcohol Depend, 100, 234–239.

Wilkinson, L.S., Mittleman, G., Torres, E., Humby, T., Hall, F.S. & Robbins, T.W. (1993) Enhancement of Amphetamine-Induced Locomotor-Activity and Dopamine Release in Nucleus-Accumbens Following Excitotoxic Lesions of the Hippocampus. Behav Brain Res, 55, 143–150.

Wilson, M.A. & McNaughton, B.L. (1993) Dynamics of the hippocampal ensemble code for space. Science, 261, 1055–1058.

Wong, S.A., Thapa, R., Badenhorst, C.A., Briggs, A.R., Sawada, J.A. & Gruber, A.J. (2017) Opposing Effects of Acute and Chronic D-Amphetamine on Decision-Making in Rats. Neuroscience, 345, 218–228.

Yu, B.M., Cunningham, J.P., Santhanam, G., Ryu, S.I., Shenoy, K.V. & Sahani, M. (2009) Gaussian-Process Factor Analysis for Low-Dimensional Single-Trial Analysis of Neural Population Activity (vol 102, pg 614, 2009). J Neurophysiol, 102, 2008–2008.

